# Impairments in *SHMT2* expression or cellular folate availability reduce oxidative phosphorylation and pyruvate kinase activity

**DOI:** 10.1101/2022.04.07.487511

**Authors:** Joanna L. Fiddler, Jamie E. Blum, Luisa F. Castillo, Anna E. Thalacker-Mercer, Martha S. Field

## Abstract

**Background:** Serine hydroxymethyltransferase 2 (SHMT2) catalyzes the reversible conversion of tetrahydrofolate (THF) and serine producing THF-conjugated one-carbon units and glycine in the mitochondria. Biallelic *SHMT2* variants were identified in humans and suggested to alter the protein’s active site, potentially disrupting enzymatic function. SHMT2 expression has also been shown to decrease with aging in human fibroblasts. Immortalized cells models of total *SHMT2* loss or folate deficiency exhibit decreased oxidative capacity and impaired mitochondrial complex I assembly and protein levels, suggesting folate-mediated one-carbon metabolism (FOCM) and the oxidative phosphorylation system are functionally coordinated. This study examined the role of SHMT2 and folate availability in regulating mitochondrial function, energy metabolism, and cellular proliferative capacity in both heterozygous and homozygous cell models of reduced *SHMT2* expression.

**Methods:** Primary mouse embryonic fibroblasts (MEF) were isolated from a C57Bl/6 dam crossed with a heterozygous *Shmt2*^*+/-*^ male to generate *Shmt2*^*+/+*^ (wild-type) or *Shmt2*^*+/-*^ (HET) MEF cells. In addition, haploid chronic myeloid leukemia cells (HAP1, wild-type) or HAP1 cells lacking SHMT2 expression (ΔSHMT2) were cultured for 4 doublings in either low-folate or folate-sufficient culture media. Cells were examined for proliferation, total folate levels, mtDNA content, protein levels of pyruvate kinase and PGC1α, pyruvate kinase enzyme activity, mitochondrial membrane potential, and mitochondrial function.

**Results:** Homozygous loss of *SHMT2* in HAP1 cells impaired cellular folate accumulation and altered mitochondrial DNA content, membrane potential, and basal respiration. Formate rescued proliferation in HAP1, but not ΔSHMT2, cells cultured in low-folate medium. Pyruvate kinase activity and protein levels were impaired in ΔSHMT2 cells and in MEF cells exposed to low-folate medium. Mitochondrial biogenesis protein levels were elevated in *Shmt2*^*+/-*^ MEF cells, while mitochondrial mass was increased in both homozygous and heterozygous models of SHMT2 loss.

**Conclusions:** The results from this study indicate disrupted mitochondrial FOCM impairs mitochondrial folate accumulation and respiration, glycolytic activity, and cellular proliferation. These changes persist even after a potentially compensatory increase in mitochondrial biogenesis as a result of decreased SHMT2 levels.

## INTRODUCTION

Folate coenzymes, found within the folate-mediated one-carbon metabolism (FOCM) metabolic network, mediate the activation and transfer of one-carbon units for diverse cellular processes, including *de novo* purine and thymidylate (dTMP) biosynthesis, amino acid metabolism, and methionine regeneration [1,2]. The transfer of one-carbon units is compartmentalized within the nucleus, cytosol, and mitochondria [3], and serine catabolism provides the majority of one-carbon units within mammalian cells [4]. Within the mitochondria, serine hydroxymethyltransferase 2 (SHMT2) catalyzes the reversible conversion of tetrahydrofolate (THF) and serine, producing THF-conjugated one-carbon units and glycine [5]. In this pathway, the majority of serine is converted to formate, which exits the mitochondria and is used in the above mentioned nuclear or cytosolic FOCM processes [1,6,7]. Recently, biallelic *SHMT2* variants were identified in humans and suggested to alter the protein’s active site [8]. Indeed, patient fibroblasts from individuals with *SHMT2* variants display a decreased ratio of glycine to serine, suggesting disrupted enzymatic function, though the consequences of reduced activity are relatively uncharacterized.

Immortalized/transformed cell models of total *SHMT2* loss have been utilized to study the implications of impaired mitochondrial FOCM. Many of these models have examined the drivers of proliferative capacity in cancer cells, which exhibit higher levels of SHMT2 than noncancer tissue [9,10]. Interestingly, changes in SHMT2 levels have also been associated with aging; aged human fibroblasts exhibit reduced *SHMT2* expression corresponding with reduced oxygen consumption [11], suggesting SHMT2 not only plays a role in cell proliferation but also in mitochondrial energy metabolism.

Serine-derived one-carbon units can also be used for *de novo* dTMP biosynthesis within the mitochondria [5,12]. Mitochondrial DNA (mtDNA) encodes 13 proteins that are required for ATP synthesis as well as mitochondrial tRNA and rRNA molecules, and it has been recognized for decades that mutations in mtDNA increase with age, lead to diseases, and occur ∼100-fold more frequently than in the nuclear genome [13–15]. Furthermore, reduced *Shmt2* expression and folate deficiency resulted in increased uracil misincorporation in mtDNA, in a heterozygous mouse model of *Shmt2* loss, without affecting uracil misincorporation in the nuclear genome [16]. These findings are consistent with evidence that mtDNA is more sensitive to genomic instability than nuclear DNA [17].

Numerous *in vitro* studies have focused on the consequences of homozygous loss of *SHMT2* in immortalized/transformed cells, however, *in vivo* models of homozygous SHMT2 loss are embryonically lethal [16,18]. *SHMT2* expression also declines with age in human fibroblast cells [11]. Since SHMT2 is at the intersection of aging and cellular proliferation/mitochondrial metabolism, we developed a mouse embryonic fibroblast (MEF) cell model of heterozygous *Shmt2* loss to more closely mimic conditions with reduced SHMT2. Here, we describe the role of SHMT2 and folate availability in regulating energy metabolism and cellular proliferative capacity in heterozygous and homozygous cell models of *SHMT2* expression.

## METHODS

### Cell culture conditions

HAP1 cells (wild-type) and SHMT2 knockout HAP1 (ΔSHMT2) cells were obtained from Horizon Discovery: ΔSHMT2 cells were generated by Horizon Discovery using CRISPR/Cas9 and contain a 2-bp deletion in SHMT2 coding exon 2. Cells were regularly passaged in Iscove’s Modification of DMEM (IMDM; Corning) supplemented with 10% FBS and 1% penicillin/streptomycin. Because metabolism related phenotypes often do not manifest based on nutrient availability, we cultured HAP1 and ΔSHMT2 cells in 25 nM (folate-sufficient) and 0 nM (folate-deficient) folate supplemented modified DMEM medium (Hyclone; formulated to lack glucose, glutamine, B vitamins, methionine, glycine and serine) containing 10% fetal bovine serum, 1% penicillin/streptomycin, 4.5 g/L glucose, 3 g/L sodium bicarbonate, 4 nM glutamine, 200 µM methionine, 4 mg/L pyridoxine, 30 mg/L glycine, and 25 or 0 nM (6S)-5-formyl-THF.

Mouse embryonic fibroblasts were isolated from C57Bl/6J female mice bred to *Shmt2*^*+/-*^ male mice as previously described [16]. All experiments include wild-type *Shmt2*^*+/+*^ and heterozygous *Shmt2*^*+/-*^ MEF cells. Cells were regularly passaged in alpha-minimal essential medium (alpha-MEM; Hyclone Laboratories) supplemented with 10% FBS and 1% penicillin/streptomycin. For 25 and 0 nM folate supplemented experimental conditions, cells were cultured in modified alpha-MEM (HyClone; lacking glycine, serine, methionine, B vitamins, and nucleosides); modified alpha-MEM was supplemented with 10% dialyzed FBS, 200 µM methionine, 1 mg/L pyridoxine, and 25 or 0 nM (6S)-5-formyl-THF.

### Cell proliferation

HAP1 cells and ΔSHMT2 cells were seeded at 1000 cells per well in 96-well plates in 25 nM and 0 nM folate supplemented medium with the addition of 0- or 2-mM formate. The number of total and dead cells was determined at specified time points by co-staining cells with Hoechst 33342 (Life Technologies) and propidium iodide (Thermo Fisher Scientific), respectively. Cells were visualized and quantified using a Celigo imaging cytometer (Nexcelom) following the manufacturer’s instructions. The number of live cells was determined by subtracting the number of propidium iodide-positive cells from the Hoechst 33342-positive cells. Data are shown as cell proliferation normalized to cell number on day 1.

### Immunoblotting

Total protein was extracted following tissue lysis by sonication in lysis buffer (150 mM NaCl, 5 mM EDTA pH8, 1% Triton X-100, 10 mM Tris-Cl, 5 mM dithiothreitol and protease inhibitor) and quantified by the Lowry-Bensadoun assay [19]. Proteins were denatured by heating with 6X Laemelli buffer for 5 min at 95 °C. Samples were electrophoresed on 8-12% SDS-PAGE gels for approximately 60-70 min in SDS-PAGE running buffer and then transferred to an Immobilon-P polyvinylidene difluoride membrane (Millipore Corp.) using a MiniTransblot apparatus (Bio-Rad). Membranes were blocked in 5% (w/v) nonfat dairy milk in 1X TBS containing 0.1% Tween-20 for 1 h at room temperature. The membranes were incubated overnight in the primary antibody at 4 °C and then washed with 1X TBS containing 0.1% Tween-20 and incubated with the appropriate horseradish peroxidase–conjugated secondary antibody at 4 °C for 1 h at room temperature. The membranes were visualized with Clarity and Clarity Max ECL Western Blotting Substrates (Bio-Rad). Antibodies against SHMT2 (Cell Signaling, 1:1000), PKM1 (Cell Signaling, 1:1000), PKM2 (Cell Signaling, 1:1000), PGC1α (Cell Signaling, 1:1000), and GAPDH (Cell Signaling, 1:2000) were used. For antibody detection, a goat anti-rabbit IgG-horseradish peroxidase-conjugated secondary (Pierce) was used at a 1:15000 dilution. Membranes were imaged using FluorChem E (Protein Simple), and densitometry was performed with ImageJ (version 1.53a) using GAPDH as the control.

### Folate concentration analysis

Cell folate concentrations were quantified using the *Lactobacillus casei* microbiological assay as previously described [20]. Total folates were normalized to protein concentrations for each sample [19].

### Mitochondrial DNA content, membrane potential, and mitochondrial function

Total genomic DNA was isolated with the Roche High Pure PCR Template Preparation Kit per manufacturers’ protocol. Mitochondrial DNA copy number was determined by real-time quantitative PCR (Roche LightCycler® 480) as previously described [21], using LightCycler® 480 SYBR Green I Master (Roche) and 15 ng of DNA per reaction. Oligonucleotide primers for mouse Mito (F 5’- CTAGAAACCCCGAAACCAAA and R 5’- CCAGCTATCACCAAGCTCGT and mouse B2M (F 5’- ATGGGAAGCCCGAACATACTG and R 5’- CAGTCTCAGTGGGGGTGAAT (Integrated DNA Technologies).

The mitochondrial membrane potential was determined using JC-1 dye (Cayman) following the manufacturer’s instructions. J-aggregate (excitation/emission = 535/595 nm) and monomer (excitation/emission = 484/535 nm) fluorescence was measured with a SpectraMax M3 (Molecular Devices).

The mitochondrial function was measured using a Seahorse XFe24 Extracellular Flux Analyzer (Agilent Technologies). Cells were cultured in the experimental 25 and 0 nM folate conditions for 4 doublings, then seeded in the same medium and allowed to adhere for 24 hours. Basal respiration, ATP production, and extracellular acidification rate were determined following the manufacturer’s instructions for the Cell Mitochondrial Stress Test (Agilent Technologies) and normalized to total cell count.

### Pyruvate kinase enzyme activity

HAP1 cells, ΔSHMT2 cells, and MEF cells were washed with 1 X PBS, pH 7.4, and then incubated with lysis buffer (50 mM Tris-HCl, pH 7.5, 1 mM EDTA, 150 mM NaCl, 1 mM DTT, protease inhibitor cocktail) for 15 minutes on ice to lyse cells. Protein was quantified by a BCA protein assay (Pierce) and 8 ug of fresh cell lysate was loaded per reaction following the instructions of the pyruvate kinase enzyme activity assay [22]. Optical absorbance of the reaction at 340 nm was measured every 15 sec for 10 min with a SpectraMax M3 (Molecular Devices).

### Mitochondrial mass and NAD/NADH ratio

The mitochondrial mass was determined using a Citrate Synthase Activity Assay Kit (Sigma-Aldrich) following the manufacturer’s instructions. Number of mitochondria was normalized to the total protein concentration. The NAD/NADH ratio was determined using a NAD/NADH-Glo Assay (Promega) according to the manufacturer’s instructions. NAD/NADH ratio was normalized to total cell number determined by Hoechst 33342 (Life Technologies) staining as described above.

### Statistical analyses

JMP® Pro statistical software version 15 (SAS Institute Inc.) was used for all statistical analyses. Linear mixed effects models with main effects of media, genotype, and time (with time as a continuous variable), and all 2- and 3-way interactions were used to determine HAP1 and ΔSHMT2 cell proliferation. For analyses in which HAP1 and ΔSHMT2 cells or *Shmt2*^*+/+*^ and *Shmt2*^*+/-*^ MEF cells were cultured in 25 or 0 nM folate supplemented medium, results were analyzed by two-way ANOVA with Tukey post-hoc analysis to determine genotype by medium interaction and main effects of medium and genotype. For analyses in which HAP1 and ΔSHMT2 cells or *Shmt2*^*+/+*^ and *Shmt2*^*+/-*^ MEF cells were compared, results were analyzed by Student’s *t* test. All statistics were performed at the 95% confidence level (α = 0.05) and groups were considered significantly different when *p ≤ 0*.*05*. Descriptive statistics were calculated on all variables to include means and standard deviations.

## RESULTS

### Loss of SHMT2 impairs cellular folate accumulation and alters mitochondrial DNA content, membrane potential, and basal respiration in HAP1 cells

We have previously demonstrated that heterozygous *Shmt2*^*+/-*^ MEF cells exhibited a ∼50% reduction in SHMT2 protein levels compared to *Shmt2*^*+/+*^ MEF cells, and *Shmt2*^*+/+*^ and *Shmt2*^*+/-*^ MEF cells cultured in low-folate medium had a reduction in total folates [16]. To assess effects of total loss of *SHMT2*, HAP1 cells and ΔSHMT2 cells were cultured in medium containing either 25 nM (6S)5-formylTHF (folate-sufficient medium) or 0 nM (6S)5-formylTHF (low-folate medium). The 2-bp deletion in an *SHMT2* coding exon resulted in a significant reduction in SHMT2 protein levels with no visible protein in the ΔSHMT2 cells (*p* < 0.01, Figure 1A). As expected, HAP1 cells and ΔSHMT2 cells grown in low-folate medium had impaired accumulation of folate (*p* < 0.01, Figure 1B), and there was a significant genotype by folate interaction (p < 0.05, Figure 1B). ΔSHMT2 cells grown in folate-sufficient medium exhibited a trend toward reduced folate accumulation compared to HAP1 cells grown in folate-sufficient medium (p = 0.05, Figure 1B), consistent with what was observed in *Shmt2*^*+/-*^ mouse liver [16].

**Figure 1.**
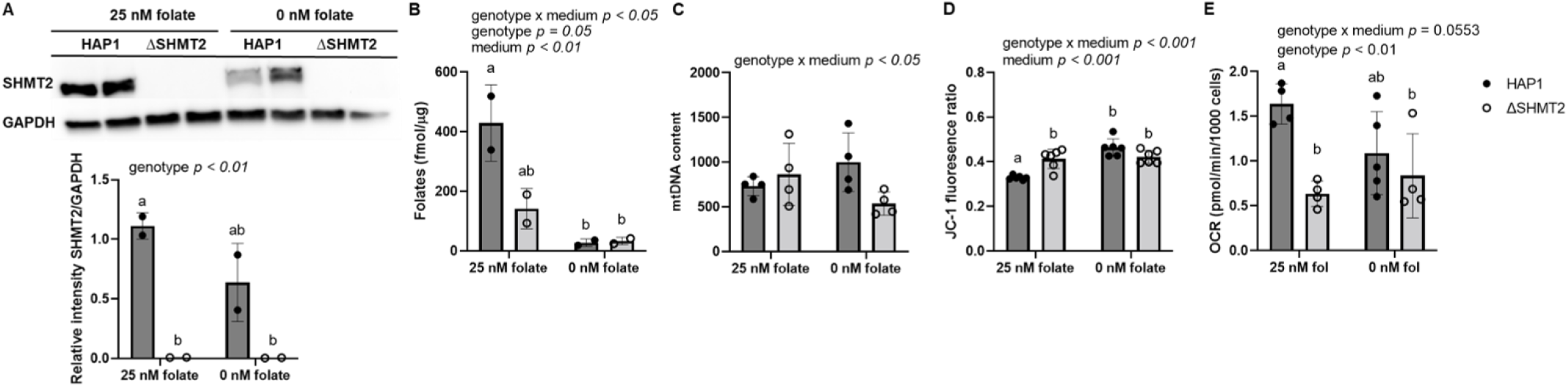
Homozygous loss of *SHMT2* and low-folate medium decrease folate accumulation and basal respiration and increase mitochondrial membrane potential in HAP1 cells. A) SHMT2 protein levels, B) total folate levels, C) mtDNA content, D) mitochondrial membrane potential, and E) oxygen consumption rate in HAP1 cells and ΔSHMT2 cells. SHMT2 protein levels were normalized to GAPDH, and densitometry was performed using ImageJ. Two-way ANOVA with Tukey’s post-hoc analysis was used to determine media by genotype interaction and main effects of media and genotype with a statistical significance at *p* < 0.05. Levels not connected by the same letter are significantly different. Data represent means ± SD values, n = 2-6 per group. GAPDH, glyceraldegyde-3 phosphate dehydrogenase; SHMT2, serine hydroxymethyltransferase 2.

There were significant genotype by medium interactions in mtDNA content (*p* < 0.05, Figure 1C) and mitochondrial membrane potential in the HAP1 and ΔSHMT2 cells (*p* < 0.001, Figure 1D). mtDNA content in HAP1 cells and ΔSHMT2 cells responded independently to folate availability, with decreased mtDNA content in ΔSHMT2 cells cultured in low-folate medium (Figure 1C). In addition, exposure to low-folate medium increased mitochondrial membrane potential in both HAP1 and ΔSHMT2 cells (*p* < 0.001, Figure 1D). Furthermore, oxygen consumption rate was impaired in ΔSHMT2 cells compared to HAP1 cells; ΔSHMT2 cells cultured in folate-sufficient medium had <50% less capacity to utilize oxygen than HAP1 cells cultured in folate-sufficient medium (*p* < 0.01, Figure 1E). A more modest ∼20% reduction in oxygen consumption was also observed in *Shmt2*^*+/-*^ MEF cells, and in *Shmt2*^*+/+*^ and *Shmt2*^*+/-*^ MEF cells cultured in low-folate medium compared to *Shmt2*^*+/+*^ MEF cells [16]. Additionally, *Shmt2*^*+/-*^ MEF cells and *Shmt2*^*+/+*^ and *Shmt2*^*+/-*^ MEF cells cultured in low-folate medium had decreased mitochondrial membrane potential with no changes in mtDNA content [16].

### The addition of formate rescues proliferation in HAP1, but not ΔSHMT2, cells cultured low-folate medium

We previously demonstrated that *Shmt2*^*+/-*^ MEF cells, and *Shmt2*^*+/+*^ and *Shmt2*^*+/-*^ MEF cells cultured in low-folate medium had reduced cellular proliferation and the addition of 2 mM formate restored cellular proliferation in *Shmt2*^*+/+*^ and *Shmt2*^*+/-*^ MEF cells cultured in low-folate medium [16]. Examination of cell proliferation rates with total loss of *SHMT2* confirmed significant effects of ΔSHMT2 genotype, exposure to low-folate media, and medium over time interaction for the cell proliferation (*p* < 0.001 for all main effects and interactions; Figure 2A). The genotype-driven differences in cell proliferation were significant at day 1 and became more pronounced at days 2 and 3 (Figure 2B). The addition of 2 mM formate rescued growth of HAP1 cells cultured in low-folate medium, but not ΔSHMT2 cells in either medium type (Figure 2A and 2C). This finding suggests that mitochondrial conversion of one-carbon units from serine to formate is not the only growth-limiting effect of *SHMT2* loss. Interestingly, 2 mM formate supplementation enhanced the proliferation capacity of HAP1 cells culture in low-folate medium compared to HAP1 cells cultured in folate-sufficient medium at day 2, but the difference was lost by day 3 and 4 (Figure 2C). ΔSHMT2 cells cultured in folate-sufficient IMDM medium had reduced cellular proliferation compared to HAP1 cells (Figure S1A-B); futhermore, the addition of formate also failed to rescue the impaired proliferation even in what is considered “complete” culture medium for HAP1 cells (Figure S1A-B).

**Figure 2.**
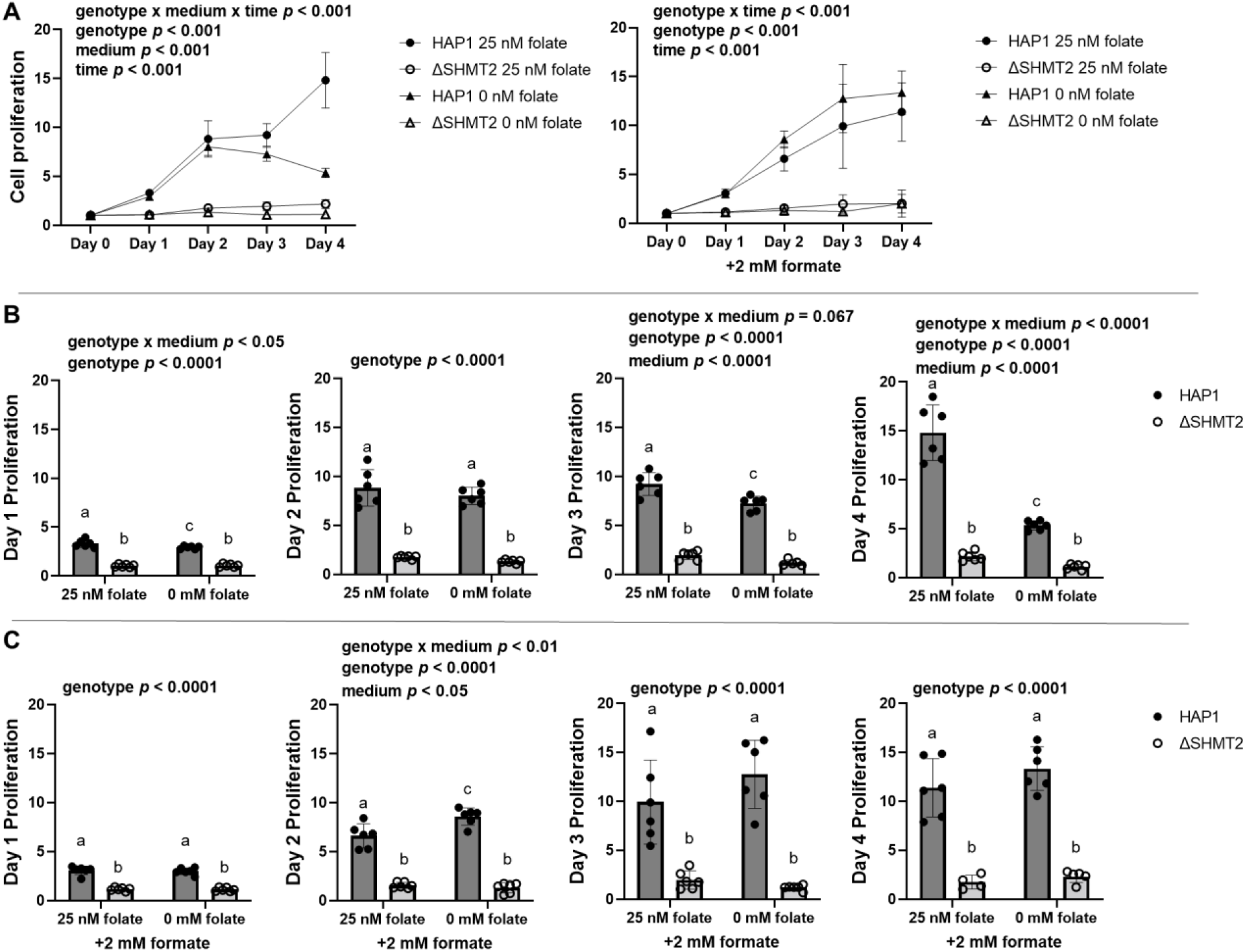
Formate rescues cell proliferation rate in HAP1 cells cultured in low-folate medium but not in ΔSHMT2 cells. Cell proliferation rates of ΔSHMT2 cells were compared with HAP1 cells by co-staining cells with Hoechst 33342 (to identify all cells) and propidium iodide (to identify dead cells). Fold change of each group was calculated by dividing by day 0 cell number. Data represent means ± SD values. Values represent n=6 replicates of cell lines cultured in medium containing either 25 nM (6S)5-formyl-THF or 0 nM (6S)5-formyl-THF. A) Cell proliferation rate and cell proliferation rate in the presence of 2 mM formate, B) relative day 1-4 quantitation of cell proliferation rate, and C) relative day 1-4 quantitation cell proliferation rate in the presence of 2 mM formate. Linear mixed effects models with main effects of media, genotype, and time (with time as a continuous variable), and 2- and 3-way interactions were used to determine cell proliferation with a statistical significance at *p* < 0.05. Two-way ANOVA with Tukey’s post-hoc analysis was used to determine media by genotype interaction and main effects of media and genotype with a statistical significance at *p* < 0.05 were used to analyze individual day proliferation. Levels not connected by the same letter are significantly different.

### Glycolytic and mitochondrial biogenesis protein levels and ATP production exhibit distinct responses in cell models of homozygous and heterozygous *SHMT2* expression

Heterozygous disruption of *Shmt2* expression in MEF cells [16] and ΔSHMT2 cells exhibit impaired oxygen consumption (Figure 1); therefore, to determine if glycolytic activity was increased to compensate for the reduction in ATP generation from oxidative phosphorylation, we assayed pyruvate kinase activity. Protein levels of PKM1 and PKM2 were significantly reduced in ΔSHMT2 cells compared to HAP1 cells (*p* < 0.01; Figure 3A). Interestingly, protein levels of PKM1 and PKM2 in *Shmt2*^*+/+*^ and *Shmt2*^*+/-*^ MEF cells were reduced only when the cells were cultured in low-folate medium, with more robust changes in PKM1 protein levels compared to PKM2 protein levels as a result of exposure to low-folate medium (Figure 4A). Pyruvate kinase enzyme activity corresponded with the proteins levels; activity was significantly reduced in ΔSHMT2 cells compared to HAP1 cells (*p* < 0.001 for genotype and genotype by medium interaction; Figure 3B), and in both *Shmt2*^*+/+*^ and *Shmt2*^*+/-*^ MEF cells cultured in low-folate medium (p < 0.001; Figure 4B). Because oxygen consumption and pyruvate kinase protein levels and enzyme activity were impaired in both cell models, we evaluated ATP production and extracellular acidification rates (ECAR). ATP production was significantly reduced in ΔSHMT2 cells compared to HAP1 cells (*p* < 0.001; Figure 5A), and in both *Shmt2*^*+/+*^ and *Shmt2*^*+/-*^ MEF cells cultured in low-folate medium (p < 0.05; Figure 5B). Interestingly, *Shmt2*^*+/-*^ MEF cells cultured in folate-sufficient medium exhibited a reduced ATP production compared to folate-sufficient *Shmt2*^*+/+*^ MEF cells (Figure 5B). Furthermore, both cell models of *SHMT2* loss and cells exposed to low-folate medium had reduced ECAR rates compared to the wild-type cells grown in folate-sufficient medium (Figure 5A and B). To determine if mitochondrial biogenesis was impacted in these models, we examined protein levels of the transcription factor PPARG Coactivator *1 Alpha* (PGC1α). *Shmt2*^*+/-*^ MEF cells cultured in folate-sufficient medium exhibited a 16-fold increase in PGC1α protein levels compared to *Shmt2*^*+/+*^ MEF cells (Figure 4A). PGC1α protein was not detected in HAP1 cells or ΔSHMT2 cells (data not shown).

**Figure 3.**
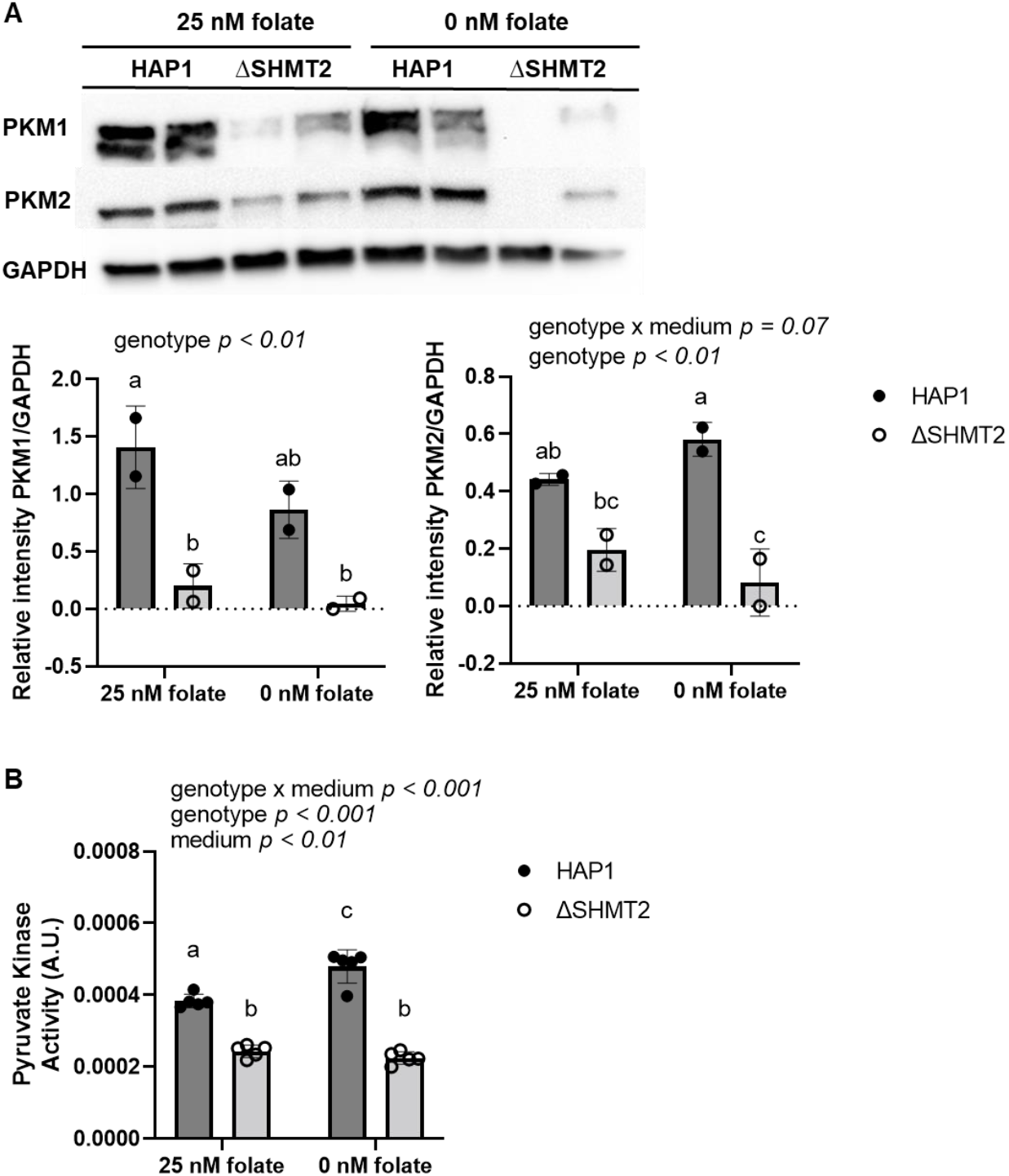
Homozygous loss of *SHMT2* reduces protein levels and activity of pyruvate kinase in HAP1 cells. A) PKM1 and PKM2 protein levels and B) pyruvate kinase activity in HAP1 cells and ΔSHMT2 cells. PKM1 and PKM2 protein levels were normalized to GAPDH and densitometry was performed using ImageJ. Two-way ANOVA with Tukey’s post-hoc analysis was used to determine media by genotype interaction and main effects of media and genotype with a statistical significance at *p* < 0.05. Levels not connected by the same letter are significantly different. Data represent means ± SD values, n = 2-5 per group. GAPDH, glyceraldegyde-3 phosphate dehydrogenase; PKM1 and PKM2, pyruvate kinase M1 and M2 isoforms.

**Figure 4.**
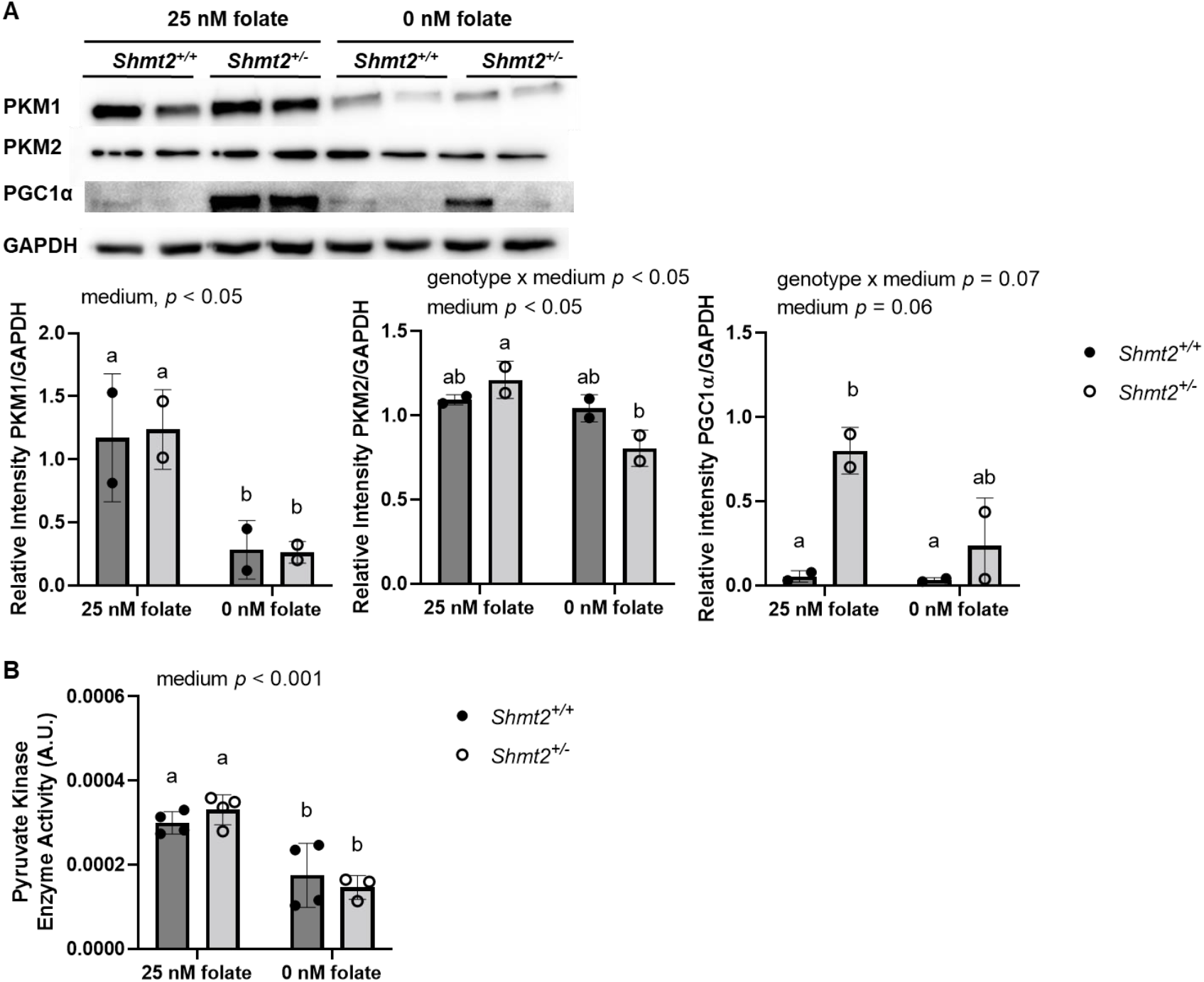
Low-folate medium decreases protein levels and activity of pyruvate kinase in MEF cells and reduced *Shmt2* expression increases PGC1α protein levels. A) PKM1, PKM2, and PGC1α protein levels and B) pyruvate kinase activity in *Shmt2*^*+/+*^ and *Shmt2*^*+/-*^ MEF cells. PKM1, PKM2, and PGC1α protein levels were normalized to GAPDH and densitometry was performed using ImageJ. Two-way ANOVA with Tukey’s post-hoc analysis was used to determine media by genotype interaction and main effects of media and genotype with a statistical significance at *p* < 0.05. Levels not connected by the same letter are significantly different. Data represent means ± SD values, n = 2-4 per group with 2 embryo cells lines represented in each group. GAPDH, glyceraldegyde-3 phosphate dehydrogenase; PGC1α, PPARγ coactivator-1α; PKM1 and PKM2, pyruvate kinase M1 and M2 isoforms.

**Figure 5.**
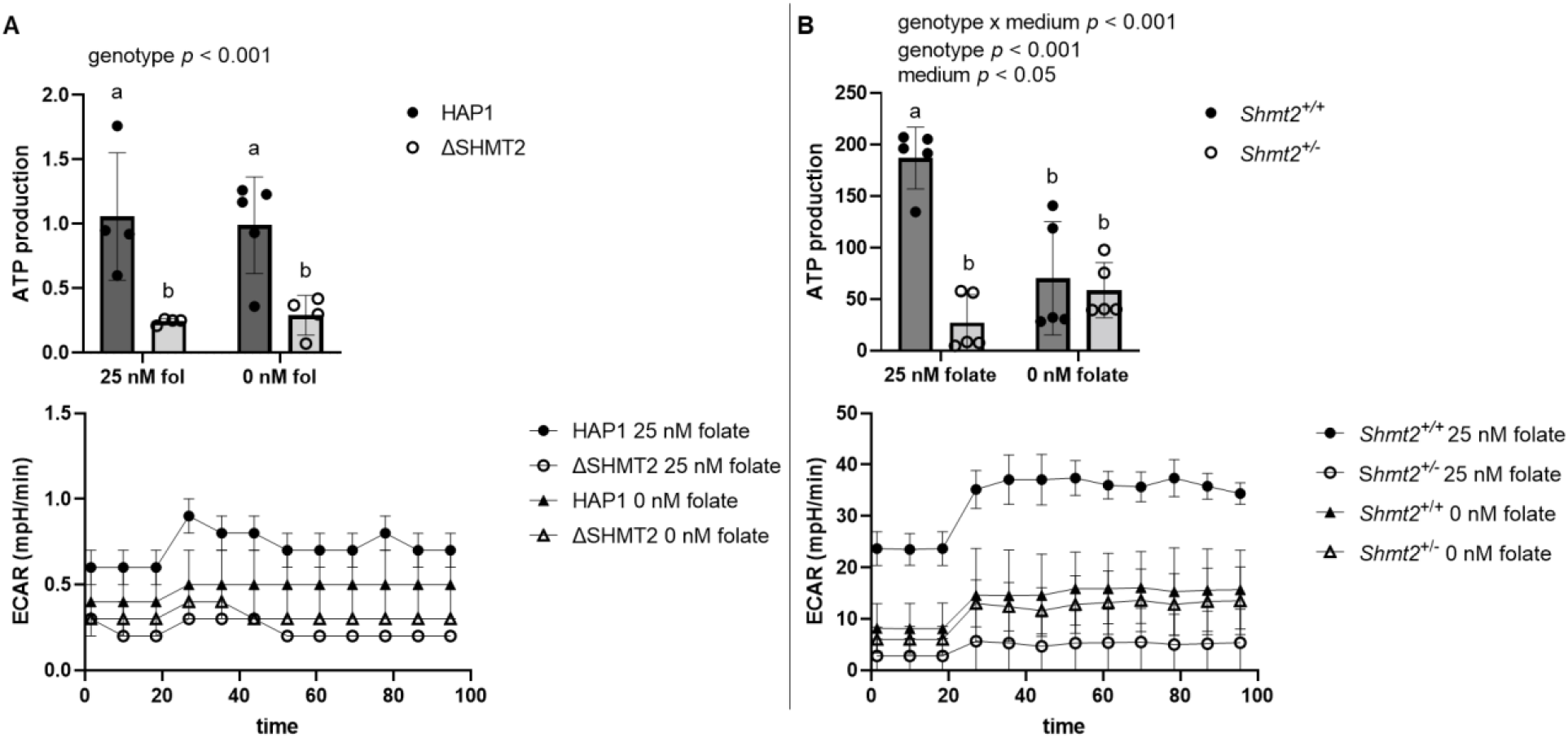
Decreased SHMT2 and exposure to low-folate medium impair ATP production and ECAR. ATP production and extracellular acidification rates in A) HAP1 cells and ΔSHMT2 cells and B) *Shmt2*^*+/+*^ and *Shmt2*^*+/-*^ MEF cells. ATP production and extracellular acidification rates were normalized to total cell count. Two-way ANOVA with Tukey’s post-hoc analysis was used to determine media by genotype interaction and main effects of media and genotype with a statistical significance at *p* < 0.05. Levels not connected by the same letter are significantly different. Data represent means ± SD values, n = 4 per group with 2 embryo cells lines represented in each group.

### Mitochondrial mass is increased as a result of homozygous and heterozygous *SHMT2* deletion, while NAD/NADH ratio is reduced only with homozygous loss of *SHMT2*

To further evaluate markers of mitochondrial health and metabolism in the homozygous and heterozygous *SHMT2* cell models, citrate synthase activity and NAD/NADH ratio were measured. Citrate synthase activity (a biomarker for mitochondrial mass [23]) was significantly increased in both ΔSHMT2 cells (p < 0.001; Figure 6A) and *Shmt2*^*+/-*^ MEF cells (p < 0.05; Figure 6B) compared to their respective wild-type cells. Additionally, there was a genotype by medium interaction in HAP1 cells and ΔSHMT2 cells cultured in low-folate and folate-sufficient medium (Figure 6A). Levels of NAD/NADH in ΔSHMT2 cells were reduced by 17% compared to HAP1 cells (p < 0.05; Figure 7A). There were no differences in NAD/NADH comparing *Shmt2*^*+/+*^ and *Shmt2*^*+/-*^ MEF cells (Figure 7B).

**Figure 6.**
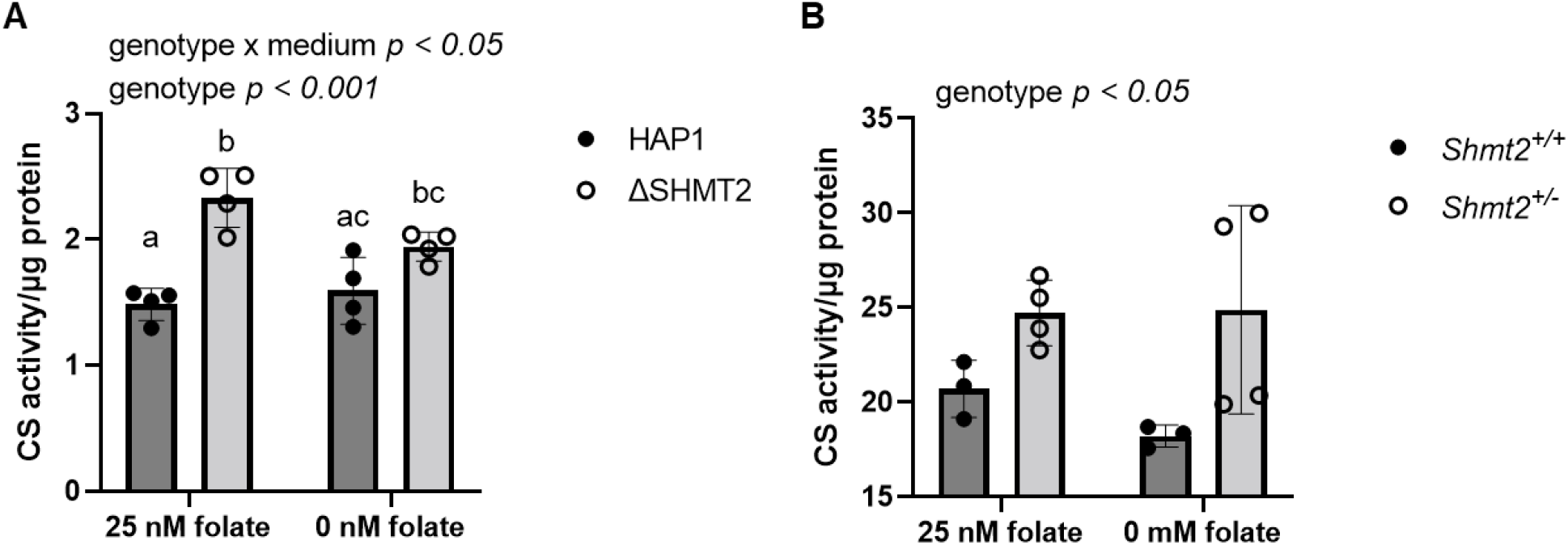
Decreased SHMT2 leads to increased mitochondrial mass. Citrate synthase activity in A) HAP1 cells and ΔSHMT2 cells and B) *Shmt2*^*+/+*^ and *Shmt2*^*+/-*^ MEF cells. Citrate synthase activity was normalized to total protein. Two-way ANOVA with Tukey’s post-hoc analysis was used to determine media by genotype interaction and main effects of media and genotype with a statistical significance at *p* < 0.05. Levels not connected by the same letter are significantly different. Data represent means ± SD values, n = 4 per group with 2 embryo cells lines represented in each group. CS, citrate synthase.

**Figure 7.**
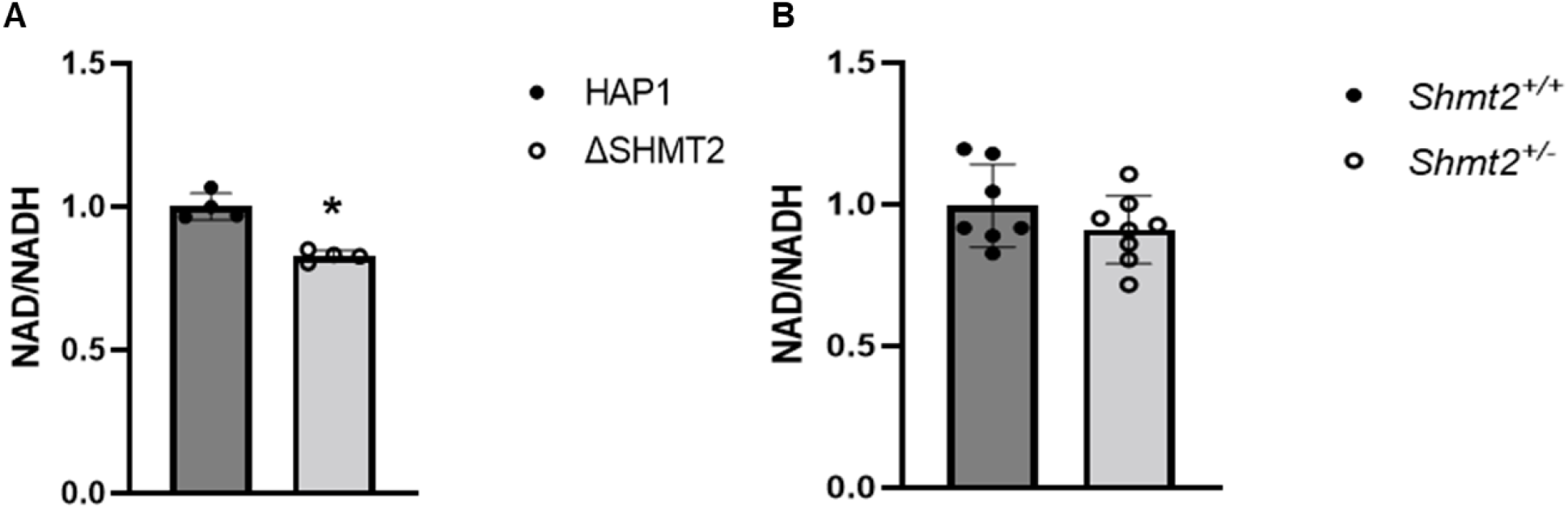
NAD/NADH ratio is impaired with homozygous *SHMT2* loss but not heterozygous loss. NAD/NADH ratio in A) HAP1 cells and ΔSHMT2 cells and B) *Shmt2*^*+/+*^ and *Shmt2*^*+/-*^ MEF cells. NAD/NADH ratio was normalized to total cell count. Two-way ANOVA with Tukey’s post-hoc analysis was used to determine media by genotype interaction and main effects of media and genotype with a statistical significance at *p* < 0.05. Levels not connected by the same letter are significantly different. Data represent means ± SD values, n = 4 per group with 2 embryo cells lines represented in each group.

## DISCUSSION

With the identification of biallelic *SHMT2* human variants [8] and the observation that *SHMT2* expression in human fibroblasts is decreased with age [11], understanding the cellular implications of perturbed mitochondrial FOCM is of great interest. In ΔSHMT2 cells, we observed decreased folate accumulation, which is consistent with our previous findings in *Shmt2*^*+/-*^ liver mitochondria [16]. Since mitochondria contain approximately 40% of total cellular folate [3], it is likely that the decreased whole-cell folate level in ΔSHMT2 cells is a result of impaired mitochondrial folate accumulation.

Effects of SHMT2 loss on cellular proliferation have varied by cell type. Multiple homozygous *SHMT2* deletion cell models indicate there are no changes in cellular proliferation rates compared to wild-type cells [24,25]. We previously demonstrated heterozygous *Shmt2* MEF cells have impaired proliferative capacity [16]. Further supporting the distinct cell type response with *SHMT2* loss; ΔSHMT2 cells cultured in either a low-folate modified medium or the more favorable IMDM medium (Figure 2 and S1) have impaired proliferative capacity compared to HAP1 cells cultured in the same media. The cell proliferative capacity and respiratory defects were not rescued by the addition of glycine in heterozygous *Shmt2* expressing MEF cells [16] or in ΔSHMT2 cells, as their growth medium contained adequate glycine levels. This suggests the one-carbon groups entering the folate pool from the glycine cleavage system are not sufficient to overcome the loss of serine-derived one-carbon units from the mitochondria to support cellular proliferation or mitochondrial function. Formate supplementation rescued HAP1 cell proliferation in cells cultured in low-folate medium (Figure 2) [16]. The inability of formate supplementation to rescue impaired proliferation in ΔSHMT2 cells suggests that the impact of SHMT2 loss on proliferation is due to effects of SHMT2 other than simply providing formate for nuclear/cytosolic FOCM, as has been suggested in other cell models [26], including in MEF cells [16].

PKM has two isoforms, PKM1 and PKM2; PKM1 is expressed in differentiated tissues (i.e., brain and muscle) while PKM2 is expressed in cancer cells, embryonic cells, and in proliferating cells [27,28]. It has previously been proposed that PKM1 promotes oxidative phosphorylation [26] and proliferation arrest due to a reduction in nucleotide biosynthesis [27], while silencing of PKM1 impairs mitochondrial membrane potential and induces apoptosis [26]. In another study, PKM2 activity is upregulated when *SHMT2* expression is suppressed [24], which suggests a lower level of SHMT2 would promote the conversion of phosphoenolpyruvate and ADP to pyruvate and ATP. In our cell models, we found that ΔSHMT2 cells had substantially reduced protein levels of PKM1 and PKM2, and reduced PK activity (Figure 3a and 3b). Independent of *Shmt2* expression levels, MEF cells exposed to low-folate medium had severely reduced protein levels of PKM1 and PK activity, with a modest reduction in PKM2 protein levels (Figure 4a and 4b). To our knowledge, this is the first observation of an effect of decreased cellular folate availability on PK activity. In addition, it does not appear that glycolysis or lactate/hydrogen ion production is increased to compensate for the reduction in mitochondrial respiration in either cell model, as both ATP production and ECAR were decreased with loss of SHMT2 or folate-depletion (Figure 5a and 5b). This is also supported by other cell models of SHMT2 loss that indicate the rate-limiting glycolytic proteins, hexokinase and phosphofructokinase, are unchanged with loss of SHMT2 [25]; however, in this cell model (293A cells), homozygous loss of SHMT2 increases lactate levels and lactate dehydrogenase protein levels. In addition to the reduction in glycolytic and respiratory capacity, it has also been demonstrated that the TCA cycle intermediates citrate, succinate, malate, and aspartate are lower in cells with homozygous loss of SHMT2 [29]. The reduced ATP production may influence the impaired proliferative capacity exhibited in ΔSHMT2 cells, *Shmt2*^*+/-*^ MEF cells, and MEF cells exposed to low-folate medium. Of note, cell health after exposure to low-folate medium indicated low cell death rates (<5%, data not shown).

Immortalized/transformed and MEF cell models of homozygous loss of *SHMT2* indicate mitochondrial derived protein levels and respiratory capacity are severely impaired, however, there were no changes in mRNA levels of these genes [18,25,30]. This supports the findings from immortalized/transformed cells models of total *SHMT2* loss that suggest reduced respiratory capacity results from impaired mitochondrial translation [29,31]. Interestingly, both ΔSHMT2 cells and *Shmt2*^*+/-*^ MEF cells have increased mitochondrial mass (Figure 6a and 6b), suggesting increased mitochondrial number in an attempt to compensate for reduced mitochondrial function. The inability to correct the impaired mitochondrial function could be from uracil misincorporation in mtDNA, as exhibited in *Shmt2*^*+/-*^ mice or *Shmt2*^*+/+*^ mice exposed to low-folate diet for 7 weeks [16], potentially causing genomic instability. The mitochondrial mass in both cell types was increased in response to loss of SHMT2, however the overall mitochondrial mass in HAP1 derived cells was much lower than in MEF cells. Another compensatory response that was only seen in *Shmt2*^*+/-*^ MEF cells was increased PGC1α protein levels. The robust increase in PGC1α protein levels in *Shmt2*^*+/-*^ MEF cells is consistent with increased mitochondrial biogenesis (i.e., increased mitochondrial mass, Figure 6b).

Regardless of whether loss of SHMT2 or folate-depletion influence mitochondrial protein translation or nucleotide biosynthesis, these findings support the notion that SHMT2 and adequate folate are essential for mitochondrial function. This may have important implications for older adults, as *SHMT2* expression in human fibroblasts declines with age [11]. Furthermore, because our cell models and that of others yield different responses, this indicates there may be tissue-specific responses to loss of SHMT2. Taken together, understanding age-associated changes in *SHMT2* expression levels, and investigating tissue-specific changes in response to reduced *SHMT2* should be assessed in future work.

## CONCLUSIONS

In this study, heterozygous and homozygous cell models of *SHMT2* expression exposed to low or adequate levels of folate were investigated. The results demonstrate that disrupted mitochondrial FOCM impairs mitochondrial folate accumulation and respiration, pyruvate kinase activity, and cellular proliferation. These findings provide evidence for the essentiality of SHMT2 and folate in maintaining energy production and have important implications for individuals with SHMT2 variants and in aging individuals.

## Abbreviations

dTMP: thymidine monophosphate
FD: folate-deficient
FOCM: folate-mediated one-carbon metabolism
GAPDH: glyceraldegyde-3 phosphate dehydrogenase
IMDM: Iscove’s modification of DMEM
MEF: murine embryonic fibroblast
MEM: minimal essential medium
mtDNA: mitochondrial DNA
PKM1: pyruvate kinase M1
OCR: oxygen consumption rate
PKM2: pyruvate kinase M2
PGC1α: PPARγ coactivator-1α
SHMT2: serine hydroxymethyltransferase 2
THF: tetrahydrofolate

## Declarations

Animal ethics approval: All mice were maintained under specific pathogen-free conditions in accordance with standard of use protocols and animal welfare regulations. All study protocols were approved by the Institutional Animal Care and Use Committee of Cornell University.

## Consent for publication

NA

## Availability of data and materials

All data generated or analyzed during this study are included in this published article [and its supplementary information files].

## Competing interests

The authors declare that they have no conflicts of interest with the contents of this article.

## Funding

This study was financially supported by a President’s Council of Cornell Women Award. Supported by National Science Foundation Graduate Research Fellowship Program grant DGE-1650441 (to JEB).

## Authors’ contributions

JLF and MSF designed research; JLF, LFC, and JEB conducted research; JLF and MSF analyzed data; JLF and MSF prepared the manuscript; and MSF has primary responsibility for the final content. All authors read and approved the final manuscript.

## Acknowledgements

NA

## Notes

### Competing Interest Statement

The authors have declared no competing interest.

