## Supplementary data for "Impairments in *SHMT2* expression or cellular folate availability reduce oxidative phosphorylation and pyruvate kinase activity"

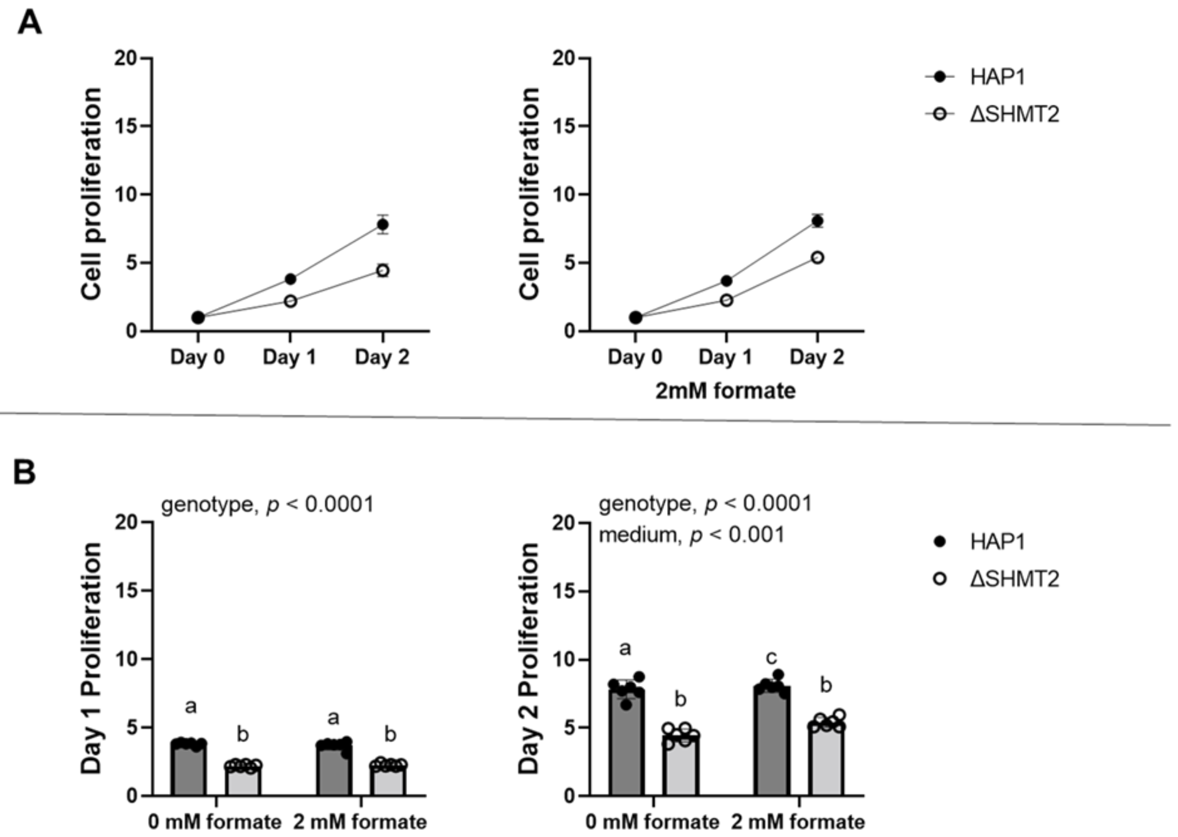


**Figure S1 Formate supplemented medium has no impact on rescuing the impaired cellular proliferation in ΔSHMT2 cells.**

Cell proliferation rates of ΔSHMT2 cells were compared with HAP1 cells by co-staining cells with Hoechst 33342 (to identify all cells) and propidium iodide (to identify dead cells). Fold change of each group was calculated by dividing by day 0 cell number. Data represent means ± SD values. Values represent n = 6 replicates of cell lines cultured in folate-sufficient IMDM medium. A) Cell proliferation rate and cell proliferation rate in the presence of 2 mM formate and B) relative day quantitation cell proliferation rate in the presence of 2 mM formate. Linear mixed effects models with main effects of media, genotype, and time (with time as a continuous variable), and 2- and 3-way interactions were used to determine cell proliferation with a statistical significance at $p < 0.05$. Two-way ANOVA with Tukey's post-hoc analysis was used to determine media by genotype interaction and main effects of media and genotype with a statistical significance at $p < 0.05$ were used to analyze individual day proliferation. Levels not connected by the same letter are significantly different.
